# Diverse Mechanisms of Resistance in Carbapenem-Resistant Enterobacteriaceae at a Health Care System in Silicon Valley, California

**DOI:** 10.1101/298513

**Authors:** Fiona Senchyna, Rajiv Gaur, Johanna Sandlund, Cynthia Truong, Guillaume Tremintin, Dietmar Küeltz, Carlos A. Gomez, Fiona B. Tamburini, Tessa Andermann, Ami Bhatt, Isabella Tickler, Nancy Watz, Indre Budvytiene, Gongyi Shi, Fred C. Tenover, Niaz Banaei

## Abstract

Carbapenem-resistant Enterobacteriaceae (CRE) are emerging as a major health threat in North America. The mechanism of resistance to carbapenems has therapeutic and public health implications. We comprehensively characterized the underlying mechanisms of carbapenem resistance in CRE isolates recovered between 2013 and 2016 at a health system in Northern California. Genotypic methods were used to detect carbapenemases and plasmid-encoded cephalosporinases, and mass spectrometry was used to quantify relative porin levels for OmpC and OmpF and their analogs. MICs for imipenem-relebactam, meropenem-vaborbactam, ceftazidime-avibactam, and ceftolozane-tazobactam were measured. Whole genome sequencing was used for strain typing. A carbapenemase gene encoding *bla*_OXA-48 like_, *bla*_NDM_, *bla*_KPC_, *bla*_SME_, *bla*_IMP_, and *bla*_VIM_ was detected in 38.7% (24/62) of CRE isolates. Porin levels was down at least 2-fold in 91.9% (57/62) of isolates. Including carbapenemase genes and porin loss, the mechanism of resistance was identified in 95.2% (59/62) of CRE isolates. Of the carbapenemase gene-positive isolates, *bla*_KPC_ -positive isolates were 100% susceptible to ceftazidime-avibactam, meropenem-vaborbactam, and imipenem-relebactam; *bla*_OXA-48 like_-positive isolates were 100% susceptible to ceftazidime-avibactam; and *bla*_SME_-positive isolates were 100% susceptible to meropenem-vaborbactam and ceftolozane-tazobactam. 100% (38/38), 92.1% (35/38), 89.5% (34/38), and 31.6% (12/38) of carbapenemase gene-negative CRE isolates were susceptible to ceftazidime-avibactam, meropenem-vaborbactam, imipenem-relebactam, and ceftolozane-tazobactam, respectively. None of the CRE strains were genetically identical. In conclusion, at this health system in Silicon Valley, carbapenemase-producing CRE occurred sporadically and were mediated by diverse mechanisms. Nucleic acid testing for *bla*_OXA-48 like_, *bla*_NDM_, *bla*_KPC_, *bla*_IMP_, and *bla*_VIM_ was sufficient to distinguish between carbapenemase-producing and non-producing CRE and accurately predicted susceptibility to ceftazidime-avibactam, meropenem-vaborbactam and imipenem-relebactam.

## Introduction

Carbapenemase-producing carbapenem-resistant *Enterobacteriaceae* (CP-CRE) have successfully spread worldwide over the recent decades (1, 2). In some regions, CP-CRE have become endemic in hospital settings (2). The proportion of CRE in acute-care hospitals in the U.S. has increased steadily (3, 4). Infection with CRE is associated with increased morbidity and mortality (5–7). Thus, early diagnosis of CRE infection is essential for timely initiation of targeted-antimicrobial therapy and early implementation of infection prevention precautions aimed to prevent nosocomial spread (8, 9).

The main mechanisms of resistance to carbapenems in CRE include hydrolysis of carbapenems by a plasmid-encoded carbapenemase, impaired outer membrane permeability due to inactivation of particular porins (i.e., OmpC and OmpF in *E. coli* and their analogs) coupled with high-level expression of cephalosporinases such as AmpC and/or extended spectrum β-lactamase (ESBL), or a combination of these mechanisms (2, 10). Moreover, CRE commonly encode genetic determinants of resistance to other classes of antibiotics rendering them pan-resistant (11). The underlying mechanism of resistance in CRE has prognostic and therapeutic implications for the newer β-lactam combination drugs that are approved by the FDA (e.g., ceftazidime-avibactam and meropenem-vaborbactam), and for those in clinical trials (e.g., imipenem-relebactam) (7, 12–14). For example, *in vitro* studies have shown the ceftazidime-avibactam combination to be effective against isolates harboring serine carbapenemases such as class A β-lactamases *Klebsiella pneumoniae* carbapenemase (KPC) and class D β-lactamases OXA-48 but not class B metallo β-lactamases (Verona integron encoded Metallo β-lactamse (VIM), IMP, or New Delhi Metallo β-lactamase (NDM)) (12). However, metallo β-lactamase-producing CRE are susceptible to avibactam when combined with aztreonam (13, 14). Thus, it is important to know both the local prevalence of CRE resistance mechanisms, and the *in vitro* susceptibility to newer β-lactam-β-lactamase inhibitor antibiotics.

Although the incidence of CRE has been lower on the West Coast of the United States compared with Eastern States (4), region-specific data are not available and comprehensive phenotypic and genotypic characterization of CRE isolates in Northern California has not been performed. We have previously reported sporadic isolation of CP-CRE at our institution (15, 16). The aim of this study was to comprehensively characterize the mechanism of carbapenem resistance in CRE isolates at our institution in Silicon Valley over a four-year period and to correlate the underlying mechanisms of resistance with susceptibility to newer β-lactam-β-lactamase inhibitor antibiotics.

## Materials and Methods

### Ethics

This study was approved by the Stanford University Internal Review Board.

### CRE isolates

All consecutive CRE isolates were included from all clinical sources from patients who received medical care at Stanford Health Care or Lucille Packard Children’s Health between January 2013 and December 2016. Both health systems consist of a tertiary academic hospital and affiliated clinics but without skilled nursing facilities. Only the first CRE isolate was included from each patient except for one patient with two strains on presentation. Isolates were identified by biochemical testing and matrix-assisted laser desorption and ionization time-of-flight mass spectrometry (MALDI-TOF) (Bruker Daltonics, Bremen, Germany). Carbapenem susceptibility testing was performed prospectively on MicroScan WalkAway plus System (Beckman Coulter, San Diego, CA) for non-urinary isolates and Vitek 2 (bioMérieux, Durham, NC) for urinary isolates. Imipenem and meropenem non-susceptible (i.e., intermediate or resistant) isolates were confirmed using disk diffusion. CRE isolates were prospectively tested for *bla*_KPC_, *bla*_NDM_, *bla*_IMP_, *bla*_VIM_ and *bla*_OXA-48 like_ using a laboratory-developed PCR assay and results were reported to providers. CRE was defined per the pre-2015 Center for Disease Control (CDC) CRE surveillance definition (17) as nonsusceptibility to imipenem (i.e., MIC > 1 μg/mL) or meropenem (i.e., MIC > 1 μg/mL), or doripenem (i.e., MIC > 1 μg/mL), and resistance to all third generation cephalosporins tested except for *Serratia marcescens* expressing *bla*_SME_, which can be susceptible to third generation cephalosporins.

### Chart review

Electronic medical records were reviewed to obtain demographics and clinical characteristics of patients with CRE infection.

### Antibiotic susceptibility testing

The following tests were performed retrospectively for research purposes. Minimum inhibitory concentrations (MIC) for imipenem, meropenem, and ertapenem were determined for urinary isolates using the MicroScan WalkAway plus System (Beckman Coulter) for comparison to non-urine isolates. Susceptibility testing for ceftolozane-tazobactam and ceftazidime-avibactam was performed by Etest (bioMérieux, Durham, NC) and meropenem-vaborbactam was performed by MIC test strip (Liofilchem Diagnostici, Teramo, Italy). Imipenem-relebactam was tested with microbroth dilution method. Relebactam was tested at a fixed concentration of 4 μg/mL. Interpretation of antimicrobial susceptibility testing results was done according to Clinical and Laboratory Standards Institute (CLSI) criteria (18) except for imipenem-relebactam and meropenem-vaborbactam which were interpreted using imipenem CLSI MIC breakpoints and package insert, respectively (18).

### Genotypic β -lactamase testing

Isolates were screened for plasmid-encoded ESBL and AmpC cephalosporinases using the Check-Points CT 103 XL Check-MDR assay (Wageningen, The Netherlands) per the package instruction. The Check-Points assay detects the following ESBLs: bla_CTX-M-1 group_, *bla*_CTX-M-1-like_, *bla*_CTX-M-15-like_, *bla*_CTX-M-32-like_, *bla*_CTX-M-2 group_, *bla*_CTX-M-8, &-25 group_, *bla*_CTX-M-9 group_, *bla*_TEM-types_, *bla*_SHV-types_, *bla*_VEB_, *bla*_PER_, *bla*_BEL_, *bla*_GES_; and the following AmpCs: *bla*_CMY I/MOX_, *bla*_ACC_, *bla*_DHA_, *bla*_ACT/MIR_, *bla*_CMY II_, *bla*_FOX_. Detection of carbapenemase genes was carried out using the Xpert Carba-R cartridge (Cepheid, Sunnyvale, CA), which detects *bla*_KPC_, *bla*_NDM_, *bla*_IMP_, *bla*_VIM_ and *bla*_OXA-48 like_; Check-Points assay which detects additionally *bla*_OXA-23 like_, *bla*_OXA-58 like_, *bla*_SPM_, *bla*_GES_, and *bla*_GIM_; and three lab-developed multiplexed PCR assays which detect *bla*_SME_*, bla*_SIM_, *bla*_SPM_, *bla*_GES_, *bla*_IMI_, *bla*_NMC-A_, and *bla*_GIM_ (Table 1). DNA was extracted by boiling a bacterial colony in molecular-grade water for 10 min. PCR reactions consisted of 2 μL of forward and reverse primer to achieve 0.5 μM, 5 μL of 2× FastStart SYBR Green Master mix (Roche Applied Science, Indianapolis, IN), and 3 μL of DNA extract. The reactions were run on a Rotor-Gene 6000 real-time cycler (Qiagen, Germantown, MD) with following cycling parameters: 95°C for 5 min and 40 cycles of 95°C for 15 sec, 60°C for 30 sec, and 72°C for 30 sec, followed by melting with ramping from 60°C to 95°C in 0.2°C increments. Melting curve analysis was performed to identify the amplicons (Table S1). Positive controls for each carbapenemase included *bla*_SME_-positive *S. marcescen*s MBRL055 and *bla*_IMI_-positive *Enterobacter cloacae* MBRL1077 provided by the Mayo Clinic (Rochester, MN); *bla*_SIM_-positive *Acinetobacter baumannii* YMC 03/9/T104 provided by Yonsei University College of Medicine (Seoul, South Korea); *bla*_GIM_-positive *E. cloacae* M15 provided by Heinrich Heine University Düsseldorf (Düsseldorf, Germany); *bla*_NMC-A_-positive *E. cloacae* and *bla*_GES-_positive *A. baumannii* provided by JMI Laboratories (North Liberty, IA); and 5 *bla*_GES-_positive and 5 *bla*_SPM-_positive *Pseudomonas aeruginosa* isolates provided by Merck (Schaumburg, IL).

**Table 1.** Annual CRE rates at Stanford Health Care

| Species | No. of CRE/CRE + non-CRE isolates (%) |  |  |  |  |
| --- | --- | --- | --- | --- | --- |
|  | 2013 | 2014 | 2015 | 2016 | 2013-16 |
| *Citrobacter freundii* complex | 0/120 (0) | 2/81 (2.5) | 0/93 (0) | 1/108 (0.9) | 3/402 (0.7) |
| *Citrobacter koseri* | 0/68 (0) | 0/54 (0) | 0/71 (0) | 0/74 (0) | 0/267 (0) |
| *Enterobacter aerogenes* | 2/122 (1.6) | 1/92 (1.1) | 4/104 (3.8) | 1/115 (0.9) | 8/433 (1.8) |
| *Enterobacter cloacae* complex | 3/240 (1.3) | 0/226 (0) | 3/239 (1.3) | 8/289 (2.8) | 14/994 (1.4) |
| *Escherichia coli* | 1/3117 (0) | 4/2080 (0.2) | 2/3008 (0.1) | 4/3834 (0.1) | 11/12039 (0.1) |
| *Klebsiella oxytoca* | 0/167 (0) | 0/129 (0) | 0/127 (0) | 0/160 (0) | 0/583 (0) |
| *Klebsiella pneumoniae* | 4/631 (0.6) | 7/503 (1.4) | 7/565 (1.2) | 1/824 (0.1) | 19/2523 (0.8) |
| *Morganella morganii* | 0/58 (0) | 0/47 (0) | 0/60 (0) | 0/66 (0) | 0/231 (0) |
| *Proteus mirabilis* | 0/245 (0) | 0/178 (0) | 0/288 (0) | 0/306 (0) | 0/1017 (0) |
| *Proteus vulgaris* | 0/16 (0) | 0/13 (0) | 0/11 (0) | 0/12 (0) | 0/52 (0) |
| *Salmonella enterica* | 0/56 (0) | 0/34 (0) | 0/29 (0) | 0/37 (0) | 0/156 (0) |
| *Serratia marcescens* | 1/161 (0.6) | 0/113 (0) | 2/157 (1.3) | 4/143 (2.8) | 7/574 (1.2) |
| All species | 11/5001 (0.2) | 14/3550 (0.4) | 18/4752 (0.4) | 19/5968 (0.3) | 62/19271 (0.3) |

### Carbapenemase activity

CRE isolates were tested for carbapenemase activity using the modified carbapenem inactivation method (mCIM) as previously described (19). Isolates with indeterminate mCIM result were tested for carbapenemase activity (i.e., imipenem degradation) with MALDI-TOF (Bruker Daltonics) as previously described (20).

### Porin protein expression

Levels of OmpC and OmpF in *E. coli* and their analogs in other species were measured using mass spectrometry (MS). Isolates were cultured overnight in 20 mL of LB broth shaking at 250 revolutions per min at 37°C. Bacterial pellets were washed in sodium phosphate buffer (SPB) and resuspended in 0.5 mL of SPB and transferred to O-ring tubes containing 0.2 mL of 0.1-mm zirconia/silica beads. Bacteria were mechanically disrupted with three 0.5-min pulses at 2,500 oscillations per min in a Mini-BeadBeater-1 (BioSpec Products, Bartlesville, OK) with 1-min intervals on ice. The lysates were sedimented two times for 10 min at 1,500 × g to remove cellular debris. To enrich for membrane proteins, the supernatants were sedimented two times for 30 min each at 21,000 × g and the second pellet was resuspended in 45 μL of SPB. Protein concentrations were measured using the Quick Start™ Bradford Protein Assay (Bio Rad, Hercules, CA) and 20 μg was separated on a 10% SDS-PAGE gel. Gels were stained with Coomassie Brilliant Blue R-250 and protein bands with molecular weight between 31 and 40 kDa were cut and digested with in-gel tryptic digestion kit (Thermo Scientific, Waltham, MA) per the package insert. Samples were concentrated in thermo savant iss110 speedvac system (Thermo Scientific) and resuspended in 20 µL of 0.1% formic acid in LC-MS grade water. Tryptic peptides (2 µL for each sample) were injected with a nanoAcquity sample manager (Waters, Milford, MA), trapped for 1 min at 15 μL/min on a Symmetry trap column (Waters), and separated on a 1.7 μm particle size BEH C18 column (Waters) by reversed phase LC using a nanoAcquity binary solvent manager (Waters). A 30 min linear acetonitrile gradient (3–35%) was applied. Peptides were ionized by nano-ESI using a pico-emitter tip (New Objective, Woburn, MA) and analyzed by an Impact HD UHR-QTOF mass spectrometer (Bruker Daltonics) in data-dependent acquisition mode. The acquisition parameters and batch processing conditions used for DDA have been previously reported (21). Data was analyzed in PreView (Protein Metrics, San Carlos, CA) using the SwissProt FASTA database entries for *Enterobacteriaceae* (www.uniprot.org) to determine the dominant post-translational modifications and mass calibration parameters. A more specific search was carried out in Byonic (Protein Metrics, San Carlos, CA) using the TrEMBL database filtered for the taxonomy of the particular organism under study. MS and MS/MS tolerances were respectively set to 10 and 30 ppm. The main modifications considered were cysteine trioxidation, methionine oxidation and N-Term acetylation. The protein false detection rate was set to 1% and all matches with less than 2 unique peptides were discarded. The resulting protein lists were then compiled with an R script (http://www.R-project.org/) to classify the identified porin variants based on homology into OmpC (OmpK36 used for *K. pneumoniae*) and OmpF (OmpK35 for *K. pneumoniae*) categories. The total intensity of all the MS/MS spectra contributing to peptide identification for each category was summed. Fold change in relative porin expression was determined by calculating the ratio of each porin in CRE isolates to averaged expression in four pan-sensitive strains of the same species.

### Porin RNA expression

Porin RNA expression was performed on the 39 CRE isolates recovered between 2013 and 2015 excluding *S. marcescens* isolates and one *E. cloacae* complex. CRE isolates were cultured in Mueller Hinton broth in the presence of a carbapenem (either meropenem 2 μg/mL or imipenem 2 μg/mL and if necessary ertapenem 1μg/mL) at a starting density 1×10^5^ CFU/mL and harvested at 1×10^8^ CFU/mL. RNAprotect Bacteria Reagent (Qiagen) was added to cultures at a ratio 3:1 and incubated at ambient temperature for 5 min. RNA was extracted from bacterial pellets and DNase-treated using RNA Extraction Kit and RNase-free DNase Kit (Qiagen), respectively. cDNA was constructed using the QuantiTect Reverse Transcription Kit (Qiagen). An identical reaction not treated with reverse transcriptase was included to control for genomic DNA carryover. Quantitative reverse transcription-PCR (qRT-PCR) was performed for porin genes (*ompC* and *ompF* in *E. coli* and their analogs in other species) and the housekeeping gene *rpoB*. qRT-PCR primers are shown in Table S2. Expression profiling of *ompF* analog in *E. cloacae* and *Citrobacter freundii* was not performed due to lack of PCR primers. PCR reactions were carried out in 10 μL containing of 0.5 µM of each primer, 1× FastStart SYBR Green Master mix, and 3 μL of cDNA. Amplification conditions were as described above. The specificity of PCR products was confirmed by melting point analysis. The cDNA copy number of each gene was extrapolated from a standard curve prepared using serial 10-fold dilution of genomic DNA from the respective species. Expression of porin genes was normalized to *rpoB* in the same sample. Fold change in porin expression was determined by calculating the ratio of normalized porin expression in CRE isolates to a pan-sensitive control strain of the same species. Each experiment was performed in triplicate, and results were presented as mean value of three experiments. *E. coli* ATCC 25922, *E. cloacae* ATCC 13047, *E. aerogenes* ATCC 13048, *K. pneumoniae* ATCC13883, and *C. freundii* ATCC 8454 were used as negative controls and *K. pneumoniae* isolate #404 (22) was used as a positive control for porin down-regulation.

### Whole genome sequencing

Genomic DNA was extracted from bacterial cultures with the Gentra Puregene Yeast/Bact. Kit (Qiagen) per the manufacturer’s instructions. Dual-indexed sequencing libraries were prepared using the Nextera XT Sample Prep Kit (Illumina, San Diego CA). Libraries were subjected to 101bp paired-end sequencing on the Illumina HiSeq 4000 platform, to approximately 100× coverage per strain. Sequencing data was demultiplexed by unique indices. Read quality was assessed using FastQC v0.11.4 (23). Reads were deduplicated using SuperDeduper v1.4 with the start location in the read at 5 bp (-s 5) and length 50 (-l 50) (24). Deduplicated reads were then trimmed using TrimGalore v0.4.4, a wrapper for CutAdapt, with a minimum quality score of 30 for trimming (-q 30), minimum read length of 50 (--length 50) and the “--nextera” flag. Preprocessed reads for each isolate were aligned to the RefSeq reference genome for the respective species using the Burrows-Wheeler Aligner (BWA) v0.7.10 with default parameters. Pileup files were generated using Samtools v1.5 (25), and Varscan v2.3.9 was used to identify single nucleotide variants (SNVs) with at least 40× coverage (--min-coverage 40), 90% frequency (--min-var-freq 0.9), and base quality of at least 20 to support a base call (--min-avg-qual 20), with the strand filter parameter turned off (--strand-filter 0). (26). Varscan output was parsed with custom scripts to generate a consensus sequence for each sample, requiring at least 0.9 frequency to support a SNP or reference base call. SNVs between strain pairs were counted using custom scripts. To build phylogenetic trees, core genome positions were identified between all strains of a given species. Core genome positions are defined as genome positions where a base call can be made for each input genome. To permit multiple sequence alignment, 30,000 SNV positions from each core genome set were randomly subsampled and concatenated into a FASTA file using custom scripts. Multiple sequence alignment was performed with MUSCLE v3.8.31, and phylogenetic trees were computed from the resulting alignments using FastTree v2.1.7 (27, 28). Trees were visualized with FigTree v1.4.3 (http://tree.bio.ed.ac.uk/software/figtree/). An isolate of *E. coli* that was sequenced in two separate runs was analyzed with this pipeline and shown to yield zero SNVs, as one would expect for an identical strain. Genome sequences for CRE isolates were deposited in the NCBI BioSample database (accession numbers SAMN08623777-SAMN08623838) (https://www.ncbi.nlm.nih.gov/Traces/study/?acc=SRP133707).

### Statistical Analysis

Fisher’s exact test was used to compare differences in proportions. Statistical analysis was done with the software GraphPad Prism 5.0, San Diego, CA.

## Results

### CRE rates

Between 2013 and 2016, out of 19,271 non-duplicate *Enterobacteriaceae* cultures with antibiotic susceptibility results, 62 (0.32%) CRE isolates from 61 patients were identified (Table 1). Demographic and clinical characteristics of patients with CRE isolates are shown in Table 2. Annual CRE rates between 2013 and 2016 did not vary significantly (0.22%, 0.39%, 0.38%, and 0.32%, respectively) (Table 1). CRE species included *Klebsiella pneumoniae* (n=19), *Enterobacter cloacae complex* (n=14), *Escherichia coli* (n=11), *Enterobacter aerogenes* (n=8), *Serratia marcescens* (n=7), and *Citrobacter freundii complex* (n=3). Carbapenem MICs for CRE isolates ranged from ≤0.5 to >4 µg/mL (interquartile range [IQR], 4 to >4) for ertapenem, ≤1 to >8 µg/mL (IQR, ≤1 to >8) for imipenem, and ≤1 to >8 µg/mL (IQR, ≤1 to >8) for meropenem.

**Table 2.** Demographics and clinical characteristics of patients with CRE infection

| Characteristic | No. (%)<br>(n=61) |
| --- | --- |
| Female | 25 (41) |
| Age - Median (range) | 56 (1-86) |
| Inpatient | 34 (55.7) |
| Comorbidities |  |
| Immunosuppression | 33 (54) |
| Solid-tumor malignancy | 13 (21.3) |
| Solid organ transplant | 11 (18) |
| Hematological-malignancy | 8 (13.1) |
| HSCT | 1 (1.6) |
| HIV-AIDS | 0 (0) |
| Diabetes | 12 (19.6) |
| Congestive Heart Failure | 10 (16.4) |
| Cirrhosis | 4 (6.5) |
| Structural lung disease | 4 (6.5) |
| Major surgery within 30 days | 4 (6.5) |
| Prematurity | 1 (1.6) |
| No comorbidities | 7 (11.5) |
| Specimen source |  |
| Urinary tract | 24 (39.3) |
| Blood | 12 (19.7) |
| Intra-abdominal | 8 (13.1) |
| Lower respiratory tract | 7 (11.5) |
| Osteoarticular | 4 (6.5) |
| Biliary tract | 2 (3.3) |
| Skin-soft tissue | 2 (3.3) |
| Genital | 1 (1.6) |
| Rrespiratory sinus | 1 (1.6) |

HSCT, Hematopoietic Stem Cell Transplant

**Table 3.** Antimicrobial susceptibility of CRE isolates

| Antimicrobial Agent | No. (%) of susceptible isolates |  |  |  |  |  |  |  |  |
| --- | --- | --- | --- | --- | --- | --- | --- | --- | --- |
|  | All CRE<br>(n=62) | Non-CP-CRE<br>(n=38) | CP-CRE (n=24) |  |  |  |  |  |  |
|  |  |  | All<br>CP-CRE | *bla*<sub>OXA-48 like</sub><br>(n=6) | *bla*<sub>KPC</sub><br>(n=5) | *bla*<sub>NDM</sub><br>(n=5) | *bla*<sub>IMP</sub><br>(n=2) | *bla*<sub>VIM</sub><br>(n=1) | *bla*<sub>SME</sub><br>(n=5) |
| Carbapenems |  |  |  |  |  |  |  |  |  |
| Imipenem | 21 (33.9) | 16 (42.1) | 5 (20.8) | 3 (50) | 0 (0) | 0 (0) | 2 (100) | 0 (0) | 0 (0) |
| Meropenem | 21 (33.9) | 17 (44.7) | 4 (16.7) | 4 (66.7) | 0 (0) | 0 (0) | 0 (0) | 0 (0) | 0 (0) |
| Ertapenem | 4 (6.5) | 3 (7.9) | 1 (4.2) | 1 (16.7) | 0 (0) | 0 (0) | 0 (0) | 0 (0) | 0 (0) |
| Monobactams |  |  |  |  |  |  |  |  |  |
| Aztreonam | 8 (12.9) | 3 (7.9) | 5 (20.8) | 0 (0) | 0 (0) | 1 (20) | 0 (0) | 1 (100) | 3 (60) |
| Cephems |  |  |  |  |  |  |  |  |  |
| Ceftriaxone | 5 (8.1) | 0 (0) | 5 (20.8) | 0 (0) | 0 (0) | 0 (0) | 0 (0) | 0 (0) | 5 (100) |
| Ceftazidime | 7 (11.3) | 2 (5.3) | 5 (20.8) | 1 (16.7) | 0 (0) | 0 (0) | 0 (0) | 0 (0) | 4 (80) |
| Cefepime | 25 (40.3) | 18 (47.4) | 7 (29.2) | 1 (16.7) | 1 (20) | 0 (0) | 0 (0) | 0 (0) | 5 (100) |
| β-lactamase inhibitor combinations |  |  |  |  |  |  |  |  |  |
| Piperacillin-tazobactam | 12 (19.4) | 8 (21.1) | 4 (16.7) | 0 (0) | 0 (0) | 0 (0) | 0 (0) | 0 (0) | 4 (80) |
| Ceftolozane-tazobactam | 17 (27.4) | 12 (31.6) | 5 (20.8) | 0 (0) | 0 (0) | 0 (0) | 0 (0) | 0 (0) | 5 (100) |
| Ceftazidime-avibactam | 54 (87.1) | 38 (100) | 16 (66.7) | 6 (100) | 5 (100) | 0 (0) | 0 (0) | 0 (0) | 5 (100) |
| Imipenem-relebactam | 44 (71) | 34 (89.5) | 10 (41.7) | 3 (50) | 5 (100) | 0 (0) | 2 (100) | 0 (0) | 0 (0) |
| Meropenem-vaborbactam | 49 (79) | 35 (92.1) | 14(60) | 4 (66.7) | 5 (100) | 0 (0) | 0 (0) | 0 (0) | 5 (100) |
| Fluoroquinolones |  |  |  |  |  |  |  |  |  |
| Ciprofloxacin | 33 (53.2) | 24 (63.2) | 9 (37.5) | 0 (0) | 3 (60) | 0 (0) | 0 (0) | 1 (100) | 5 (100) |
| Levofloxacin | 35 (56.5) | 26 (68.4) | 9 (37.5) | 0 (0) | 3 (60) | 0 (0) | 0 (0) | 1 (100) | 5 (100) |
| Aminoglycosides |  |  |  |  |  |  |  |  |  |
| Gentamicin | 41 (66.1) | 33 (86.8) | 8 (33.3) | 1 (16.7) | 2 (40) | 0 (0) | 0 (0) | 0 (0) | 5 (100) |
| Tobramycin | 36 (58.1) | 27 (71.1) | 9 (37.5) | 2 (33.3) | 1 (20) | 0 (0) | 1 (50) | 0 (0) | 5 (100) |
| Amikacin | 50 (80.6) | 35 (92.11) | 15 (62.5) | 5 (83.3) | 3 (60) | 0 (0) | 1 (50) | 1 (100) | 5 (100) |
| Tigecycline | 54 (87.1) | 34 (89.5) | 20 (83.3) | 4 (66.7) | 5 (100) | 5 (100) | 0 (0) | 1 (100) | 5 (100) |
| Trimethoprim/Sulfamethoxazole | 35 (56.5) | 27 (71.1) | 8 (33.3) | 0 (0) | 1 (20) | 1 (20) | 1 (50) | 0 (0) | 5 (100) |

non-CP, non-carbapenemase producing; CP, carbapenemase producing

### Genotypic carbapenemase testing

CRE isolates were genotypically tested for previously characterized plasmid-encoded and chromosomally-encoded carbapenemases using the Xpert Carba-R cartridge, Check-Points microarray, and a lab-developed multiplexed, real-time PCR assay. A single carbapenemase gene was detected in 38.7% (24/62) of CRE isolates which consisted of *bla*_OXA-48 like_ (n=6), *bla*_NDM_ (n=5), *bla*_KPC_ (n=5), *bla*_SME_ (n=5), *bla*_IMP_ (n=2), and *bla*_VIM_ (n=1) (Figure 1 and Table S3). The bacterial species encoding carbapenemase genes included *E. coli*, *E. cloacae* complex, *K. pneumoniae* and *S. marcescens*. The remaining 61.3% (38/62) of CRE isolates were negative for *bla*_KPC_, *bla*_NDM_, *bla*_IMP_, *bla*_VIM_ and *bla*_OXA-48 like_, *bla*_SME_*, bla*_SIM_, *bla*_SPM_, *bla*_GES_, *bla*_IMI_, *bla*_NMC-A_, and *bla*_GIM_. The species in this group included *C. freundii* complex, *E. coli*, *E. cloacae* complex, *E. aerogenes*, *K. pneumoniae* and *S. marcescens*. Annual non-CP-CRE rates between 2013 and 2016 did not vary significantly (63.6%, 50.0%, 61.1%, and 68.4%, respectively). The “SPACE” organisms that are likely to carry chromosomal AmpC such as *Serratia*, *Citrobacter*, and *Enterobacter*, made up 63.2% (24/38) of CRE isolates lacking a carbapenemase gene compared with 33.3% (8/24, p=0.04) of isolates harboring a carbapenemase gene. ESBL and plasmid-encoded AmpC cephalosporinases were detected with the Check-Points assay in 75.0% (18/24) and 20.8% (5/24), respectively, of carbapenemase gene-positive and 34.2% (13/38, P=0.004) and 28.9% (11/38, p=0.6), respectively, of carbapenemase gene-negative isolates. Either an ESBL or plasmid-encoded AmpC gene was detected in 79.2% (19/24) of carbapenemase gene-positive and 60.5% (23/38, p=0.2) of carbapenemase gene-negative isolates. All 5 carbapenemase gene-positive CRE isolates without an ESBL or AmpC gene were *bla*_SME_-positive *S. marcescens* isolates. Compared with carbapenemase gene-negative CRE, a higher proportion of carbapenemase gene-positive CRE showed an elevated imipenem and meropenem MIC >8 (15.8% vs. 45.8%, p<0.02; 10.5% vs. 58.3%, p<0.001, respectively) (Figure 2).

**Figure 1.**
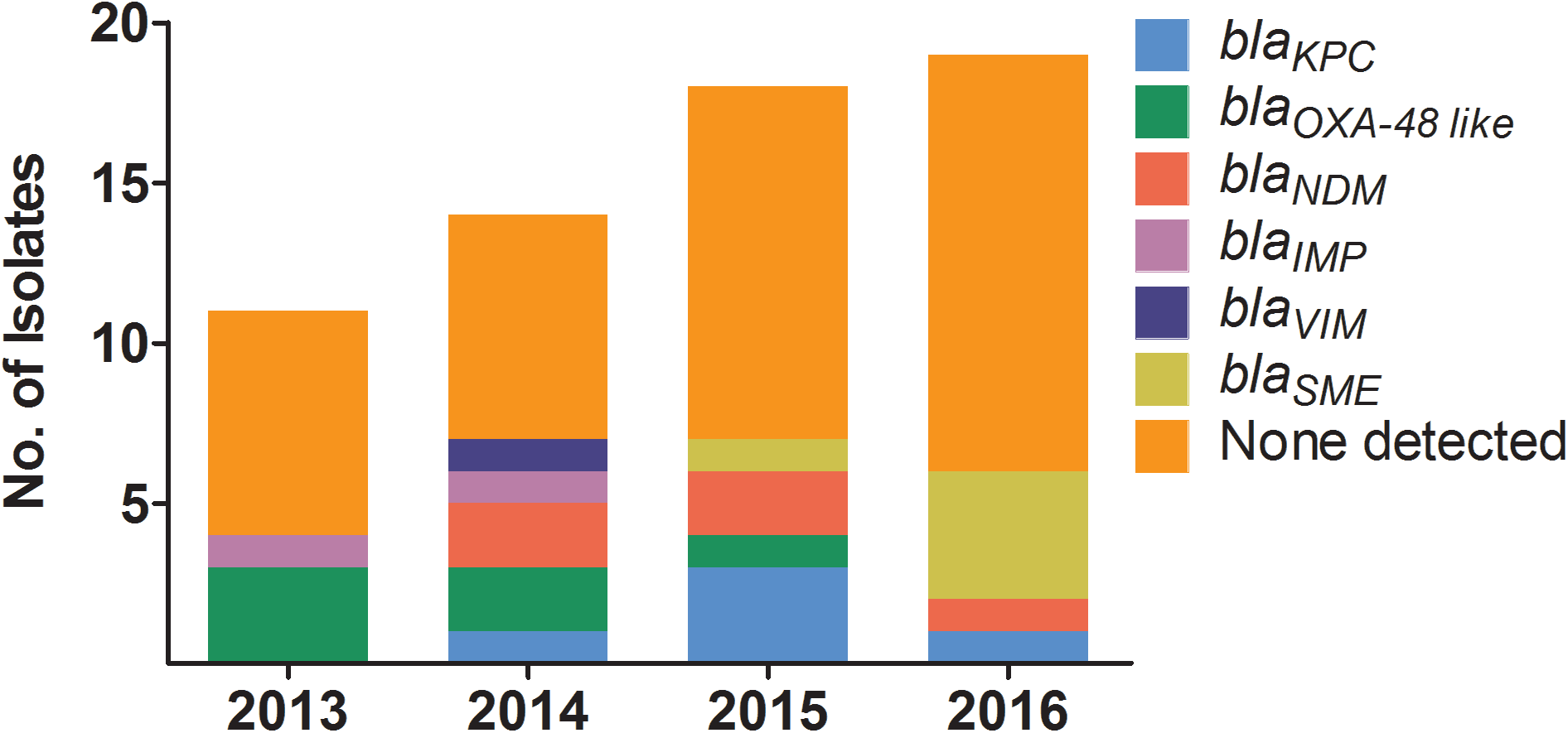
Carbapenemase genes detected in CRE isolates annually from 2013 to 2016. Carbapenemase genes tested include *bla*_KPC_, *bla*_NDM_, *bla*_IMP_, *bla*_VIM_, *bla*_OXA-48 like_, *bla*_SME_, *bla*_SIM_, *bla*_SPM_, *bla*_GES_, *bla*_IMI_, *bla*_NMC-A_, and *bla*_GIM_.

**Figure 2.**
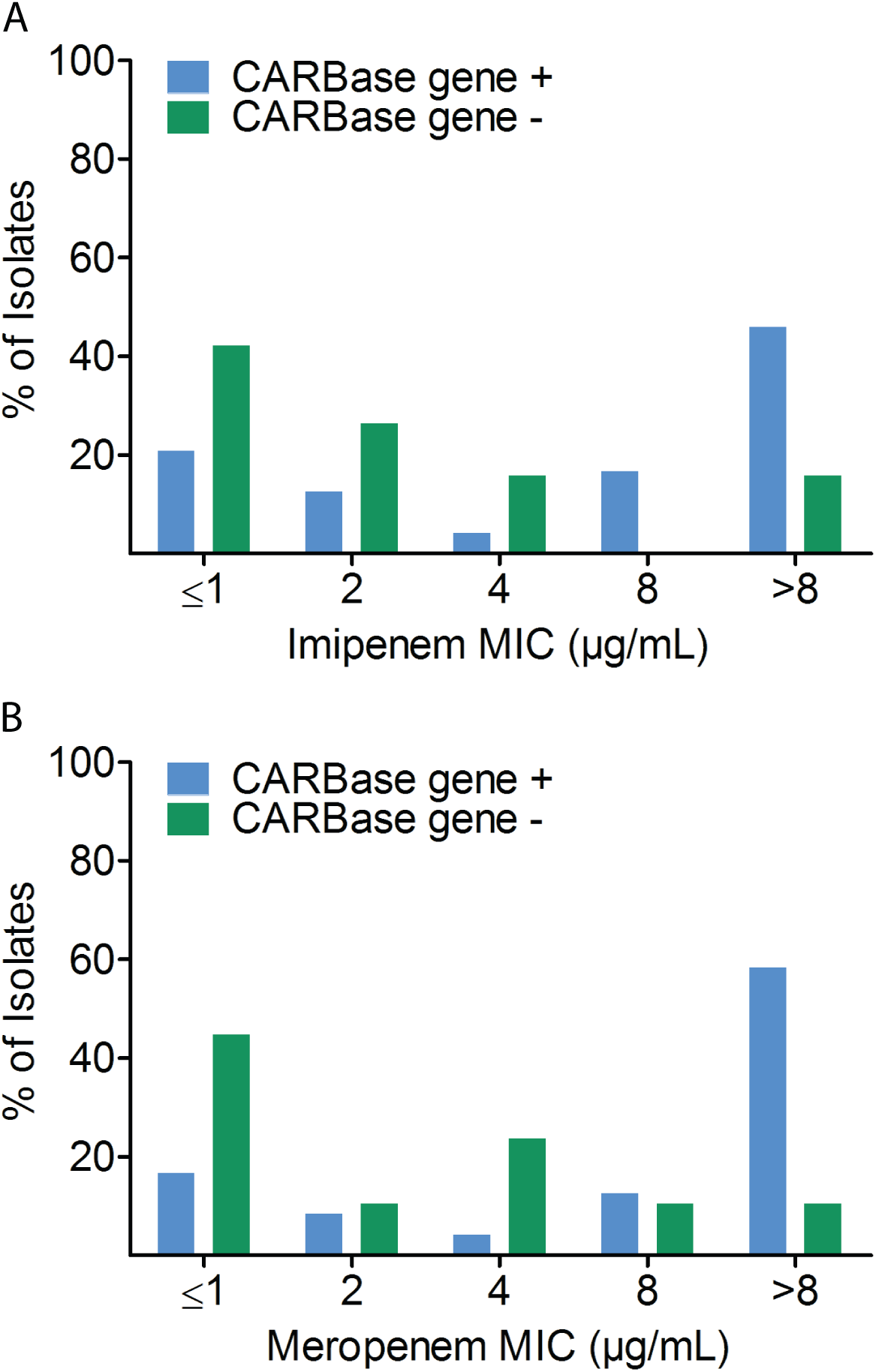
Distribution of imipenem and meropenem MICs in CRE isolates with and without a carbapenemase gene. Bars show percentage of imipenem (A) and meropenem (B) MICs for carbapenemase gene-negative (CARBase gene -; green bars) and carbapenemase gene-positive (CARBase gene +; blue bars) CRE isolates.

### Phenotypic carbapenemase testing

To determine whether carbapenemase gene-negative CRE isolates display carbapenemase activity (presumably due to previously uncharacterized carbapenemases), we performed a modified carbapenem inactivation method (mCIM) on all CRE isolates (19). While 100% (24/24) of carbapenemase gene-positive CRE isolates were mCIM-positive, 86.8% (33/38) of carbapenemase gene-negative CRE isolates were mCIM-negative, and the remaining 13.2% (5/38) were mCIM-intermediate. The mCIM-indeterminate isolates were further tested with a MALDI-TOF-based carbapenemase activity assay (20). All 5 mCIM-indeterminate isolates were confirmed carbapenemase-negative while all carbapenemase gene-positive CRE isolates evaluated tested positive for imipenem hydrolysis.

### Porin expression

To determine whether porin expression is lower in CRE isolates without a carbapenemase gene, a novel mass spectrometry-based assay was employed to measure porin proteins in all 62 CRE isolates irrespective of their species. As shown in Figure 3B, relative porin levels of OmpC and OmpF and their analogs was down 2-fold or greater in 54.2% (13/24) and 83.3% (20/24) of carbapenemase gene-positive and 71.1% (27/38, p=0.3) and 81.6% (31/38, p=1.0) of carbapenemase gene-negative CRE isolates, respectively (Figure 3B and Table S3). The expression of either OmpC or OmpF and their analogs was decreased in 91.7% (22/24) of carbapenemase gene-positive CRE isolates compared with 92.1% (35/38, p=1.0) of carbapenemase gene-negative CRE isolates. The two carbapenemase gene-positive isolates with normal porin levels were both *bla*_SME_-positive *S. marcescens* (CRE35 and 49); three carbapenemase gene-negative CRE isolates with normal porin levels were two strains of *E. cloacae complex* (CRE71 and 81), and one *C. freundii complex* (CRE21).

**Figure 3.**
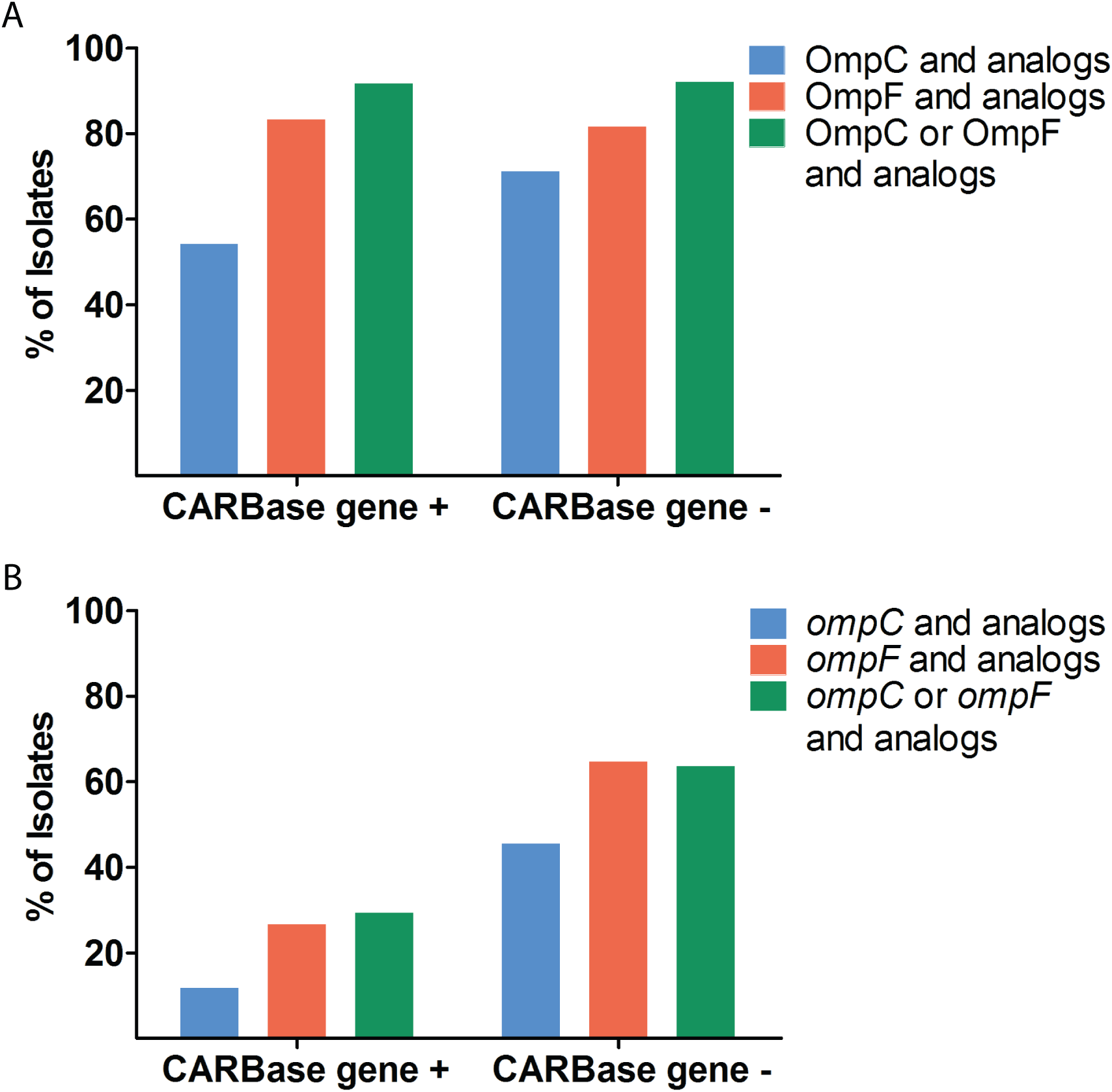
Porin protein and mRNA levels in CRE isolates with and without a carbapenemase gene. Bars show percentage of carbapenemase gene-positive (CARBase gene +) and gene-negative (CARBase gene -) CRE isolates with relative protein (A) and porin mRNA (B) down 2-fold or more compared with susceptible isolates.

We also performed quantitative reverse transcription-PCR (qRT-PCR) on 39 CRE isolates recovered between 2013 and 2015 to measure porin mRNA transcripts in CRE isolates. The expression of *ompC* and *ompF* and their analogs was downregulated 2-fold or more in 11.8% (2/17) and 26.7% (4/15) of carbapenemase gene-positive CRE isolates compared with 45.5% (10/22, p=0.04) and 64.7% (11/17, p=0.04) of carbapenemase gene-negative CRE isolates, respectively. In carbapenemase gene-negative CRE isolates, 63.6% (14/22) showed downregulation of either *ompC* or *ompF* and their analogs compared with 29.4% (5/17) in carbapenemase gene-positive CRE (p=0.05) (Figure 3A and Table S3).

### Susceptibility of CRE to newer β-lactam/β-lactamases inhibitors

In vitro studies have shown predictable susceptibility of CP-CRE to newer β-lactam/β-lactamase inhibitor combinations such as imipenem-relebactam, meropenem-vaborbactam, and ceftazidime-avibactam, depending on the molecular class of the carbapenemases they carry (12–14). We therefore investigated the susceptibility of CRE isolates to newer β-lactam/β-lactamase inhibitor combination drugs. Among carbapenemase-positive CRE isolates, 41.7% (10/24), 58.3% (14/24) and 66.7% (16/24) were susceptible to imipenem-relebactam, meropenem-vaborbactam, and ceftazidime-avibactam, respectively (Figure 4 and Table S3). Isolates that remained resistant to imipenem in the presence of relebactam were positive for class A serine β-lactamase (*bla*_SME_), class B metallo β-lactamases B (i.e., *bla*_NDM_ and *bla*_VIM_), and class D serine β-lactamase (i.e, *bla*_OXA-48 like_). Isolates that remained non-susceptible to meropenem in the presence of vaborbactam were positive for class B metallo β-lactamases B (i.e., *bla*_NDM_, *bla*_IMP_, and *bla*_VIM_) and class D serine β-lactamase (i.e, *bla*_OXA-48 like)._ All 8 ceftazidime-avibactam-resistant isolates were positive for class B metallo β-lactamases (i.e., *bla*_NDM_, *bla*_IMP_, and *bla*_VIM_)(Figure 4 and Table S3). Among carbapenemase gene-negative CRE isolates, 89.5% (34/38), 92.1% (35/38) and 100% (38/38) were susceptible to imipenem-relebactam, meropenem-vaborbactam and ceftazidime-avibactam, respectively (Figure 4 and Table S3). Three isolates that were non-susceptible to both imipenem-relebactam and meropenem-vaborbactam included *E. aerogenes* (CRE09), *E. coli* (CRE15) and *K. pneumoniae* (CRE25) (Table S3). Average relative porin levels were lower in these isolates compared with imipenem-relebactam and meropenem-vaborbactam susceptible isolates that were non-susceptible to the carbapenem alone (0.01 and 0.002 vs. 1.25 and 0.41 for OmpC and OmpF and their analogs, respectively). A *S. marcescens* CRE isolate (CRE05) was non-susceptible to imipenem-relebactam (Table S3). We also investigated the susceptibility of CRE isolates to ceftolozane-tazobactam. Among carbapenemase gene-positive CRE, 20.8% (5/24) were susceptible and 16.7% (4/24) were intermediate to ceftolozane-tazobactam (Figure 4 and Table S3). All 5 susceptible isolates were *bla*_SME_-positive *S. marcescens*. The 4 intermediate isolates consisted of a *bla*_OXA-48 like_-positive *K. pneumoniae*, a *bla*_OXA-48 like_-positive *E. coli*, a *bla*_KPC_-positive *K. pneumoniae* and a *bla*_KPC_-positive *E. cloacae complex*. Among carbapenemase gene-negative isolates, 31.6% (12/38) were susceptible and 15.8% (6/38) were intermediate to ceftolozane-tazobactam. There was no evidence of correlation between resistance to ceftolozane-tazobactam and lower porin protein levels (95.0% vs. 88.9%; p=0.3).

**Figure 4.**
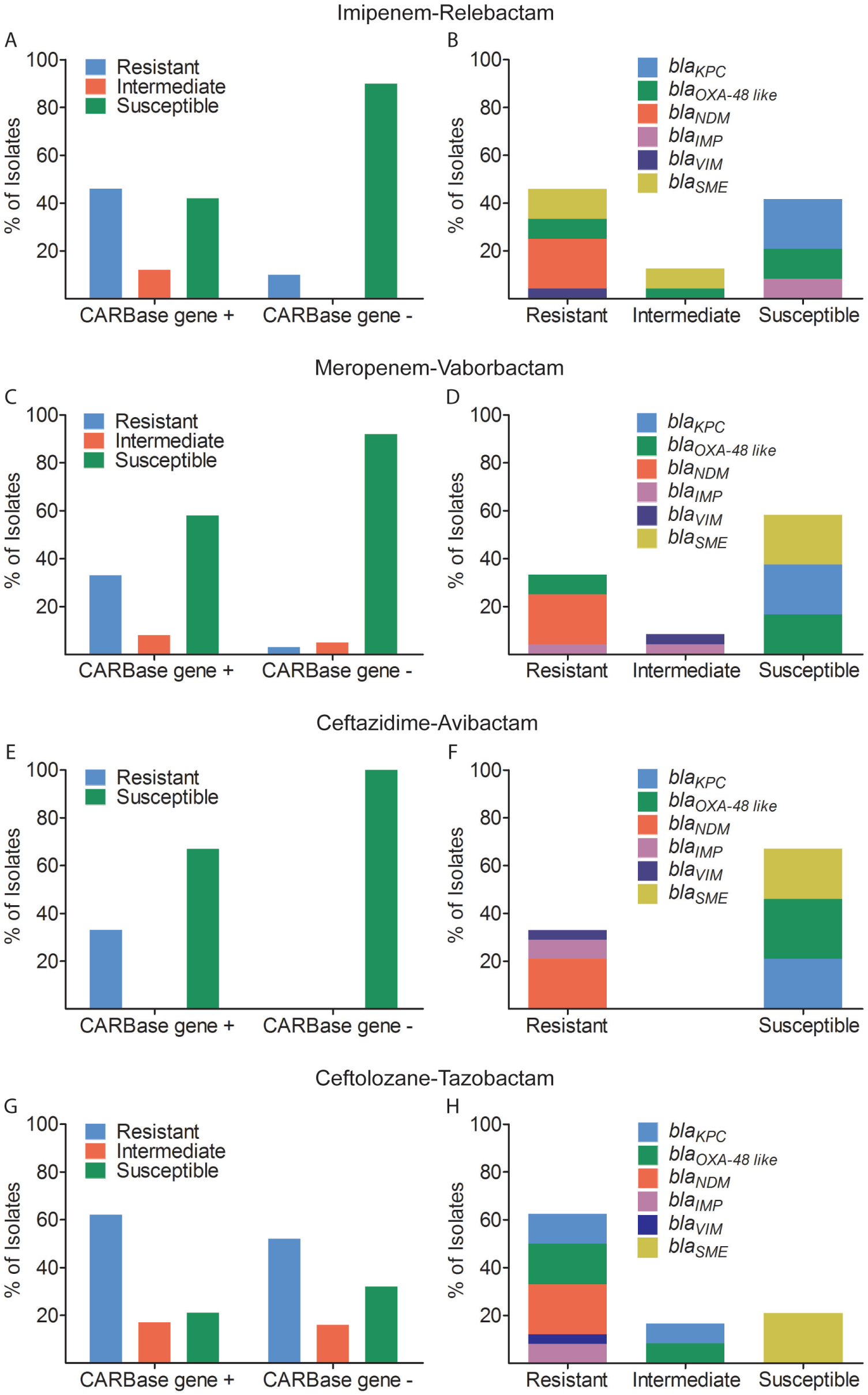
Susceptibility of CRE isolates to imipenem-relebactam, meropenem-vaborbactam, ceftazidime-avibactam, and ceftolozane-tazobactam. Bars show percent susceptibility of carbapenemase gene-positive (CARBase gene +) and gene-negative (CARBase gene -) CRE to (A) imipenem-relebactam, (C) meropenem-vaborbactam, (E) ceftazidime-avibactam, and (G) ceftolozane-tazobactam. Graphs on the right (B, D, F, H) show fraction of carbapemase genes detected in carbapenemase gene-positive isolates that are susceptible, intermediate, or resistant to the respective antibiotic combination.

### Molecular epidemiology

Whole genome sequencing was performed to investigate clonality of CRE isolates recovered between 2013 and 2016. Whole genome sequences were obtained for all CRE isolates excluding CRE30, which yielded no interpretable results. The only two CRE strains with temporal association were non-CP-CRE *K. pneumoniae* CRE24 and CRE25 which were isolated on 12/28/2014 and 12/31/2014, respectively, from patients on two different medical wards. Phylogenetic trees constructed based on whole genome sequence analysis are shown in Figure 5. We did not find any identical CRE strains among *C. freundii complex*, *E. cloacae complex*, *E. aerogenes*, *E*. *coli*, *K. pneumoniae*, and *S. marcescens* isolates (Figure 5). The number of single nucleotide variants (SNVs) for closely related strains is shown in Table 4. Phylogenetic distance between strains within a species do not indicate any transmission events, as no identical strains were observed in our data set. Even the most closely related CRE strains on the phylogenetic tree (non-CP-CRE *K. pneumoniae* CRE04 isolated on 4/16/2013 and CRE24 isolated on 12/28/2014) had 5 SNVs. The only two CRE strains with temporal association (CRE24 and CRE25) had 10 SNVs.

**Figure 5.**
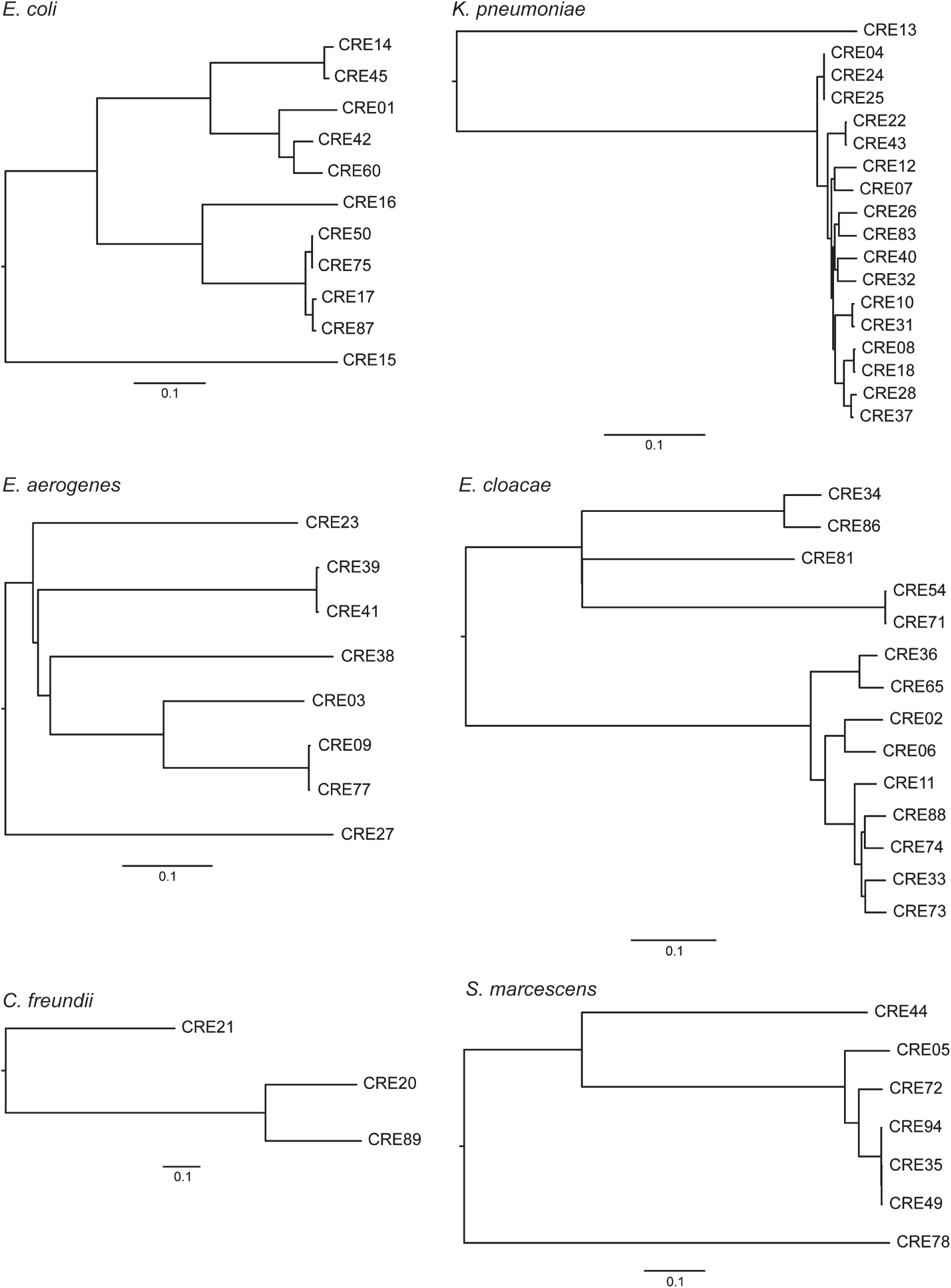
Phylogenetic tree for CRE isolates based on whole genome sequencing. Phylogenetic trees were computed from multiple sequence alignments of concatenated SNVs on a per-species basis. Scale bars show evolutionary distances.

**Table 4.** Single nucleotide variants (SNVs) of related CRE strains

| Isolate ID No. | Species | Collection Date | Carbapenemase gene | SNVs |
| --- | --- | --- | --- | --- |
| CRE50 | *E. coli* | 2/1/2016 | Negative | 136 |
| CRE75 |  | 8/16/2016 | Negative |  |
| CRE17 | *E. coli* | 10/15/2014 | *bla<sub>NDM</sub>* | 1527 |
| CRE87 |  | 11/13/2016 | *bla<sub>NDM</sub>* |  |
| CRE04 | *K. pneumoniae* | 4/6/2013 | Negative | 5 |
| CRE24 |  | 12/28/2014 | Negative |  |
| CRE04 | *K. pneumoniae* | 4/6/2013 | Negative | 7 |
| CRE25 |  | 12/31/2014 | Negative |  |
| CRE24 | *K. pneumoniae* | 12/28/2014 | Negative | 10 |
| CRE25 |  | 12/31/2014 | Negative |  |
| CRE08 | *K. pneumoniae* | 8/2/2013 | *bla<sub>OXA-48 like</sub>* | 1907 |
| CRE18 |  | 10/19/2014 | *bla<sub>NDM</sub>* |  |
| CRE22 | *K. pneumoniae* | 11/19/2014 | Negative | 163 |
| CRE43 |  | 12/24/2015 | *bla<sub>KPC</sub>* |  |
| CRE35 | *S. marcescens* | 9/2/2015 | *bla<sub>SME</sub>* | 30 |
| CRE49 |  | 1/27/2016 | *bla<sub>SME</sub>* |  |
| CRE35 | *S. marcescens* | 9/2/2015 | *bla<sub>SME</sub>* | 40 |
| CRE94 |  | 12/5/2016 | *bla<sub>SME</sub>* |  |
| CRE49 | *S. marcescens* | 1/27/2016 | *bla<sub>SME</sub>* | 40 |
| CRE94 |  | 12/5/2016 | *bla<sub>SME</sub>* |  |
| CRE39 | *E. aerogenes* | 10/30/2015 | Negative | 1258 |
| CRE41 |  | 11/18/2015 | Negative |  |
| CRE09 | *E. aerogenes* | 8/23/2013 | Negative | 176 |
| CRE77 |  | 8/30/2016 | Negative |  |
| CRE54 | *E. cloacae* | 3/1/2016 | Negative | 55 |
| CRE71 |  | 8/25/2016 | Negative |  |

## Discussion

Compared to national CRE rates of 4.2% or 1.4%, based on 2011 data submitted to National Healthcare Safety Network and 2010 data submitted to Surveillance Network-USA (4), respectively, the CRE rate of 0.32% between 2013 and 2016 at our healthcare system serving the Silicon Valley area was 4 to 13-fold lower, respectively. Although surveillance data in Northern California is lacking, our CRE rate was also lower than that reported by an academic health system in Los Angeles, California (0.32% vs. 0.73%) (29). The reason for a lower CRE rate at our institution could be due to differences in patient population (i.e., less health-care exposure), differences in infection control and prevention practices (i.e., isolation of patients with CP-CRE), absence of skilled nursing facilities and long-term-acute-care hospitals from our health system (30), and the definition used to define CRE. The second difference seems to be critical as CDC reported at least one CRE infection in 3.9% of short-stay hospitals compared with 17.8% of long-term-acute-care hospitals (4). Furthermore, in this study we applied the pre-2015 CDC CRE surveillance definition, which excludes ertapenem. Inclusion of ertapenem would have increased our CRE rate (17). While the rate of health care-associated infection caused by carbapenem-nonsusceptible *Enterobacteriaceae* has increased in the U.S between 2001 and 2011 (4), the annual CRE rates at our institution did not vary significantly over a four-year study period from 2013 to 2016.

The CP-CRE rate of 38.7% among all CRE at our institution is comparable to 47.9% reported for metropolitan areas in 7 U.S. states, including Oregon and Colorado, but much lower than 81.7% reported by an academic health system in Los Angeles (31). Unlike all other North American institutions where KPC predominates as the most common carbapenemase among CP-CRE isolates (7, 29, 31–33), at our institution CP-CRE were evenly caused by KPC (20.8%), OXA-48 like (25.0%), NDM (20.8%), and SME (20.8%), and less commonly by IMP (8.3%) and VIM (4.2%). The reason for non-predominance of KPC at our health system may be in part due to the geographic location of our institution in the Silicon Valley where high-tech industry draws people from major cities around the globe where different plasmid-encoded carbapenemases are endemic. Another important contributor is the fact that nosocomial transmission of KPC did not occur at our institution during the study period. In fact, whole genome sequence-based strain typing showed that all plasmid-encoded CP-CRE were distinct strains and therefore had occurred sporadically. As discussed above, absence of skilled nursing facilities and long-term-acute-care hospitals from our health system may have contributed to lack of CRE transmission (30).

A major strength of study is that we employed comprehensive phenotypic and genotypic analyses to determine the molecular basis of carbapenem resistance in CRE isolates recovered longitudinally at our institution. Importantly, concordance between carbapenemase activity and carbapenemase gene detection was 100%, which indicates genotypic testing would be sufficient to detect all plasmid-mediated (i.e., *bla*_KPC_, *bla*_OXA-48 like_, *bla*_NDM_, *bla*_IMP_, and *bla*_VIM_) and chromosomally-encoded (i.e., *bla*_SME_) CP-CRE at our institution. Although FDA-cleared commercial assays currently do not detect *bla*_SME_ (34), this carbapenemase should be suspected in carbapenem-resistant carbapenemase-producing *S. marcescens* isolates that are negative for *bla*_KPC_, *bla*_OXA-48 like_, *bla*_NDM_, *bla*_IMP_, and *bla*_VIM_ (35). Resistance in non-CP-CRE is mediated through high-level expression of cephalosporinases such as AmpCs and ESBLs coupled with porin inactivation (2, 10). Using a novel mass spectrometry assay, we showed porin levels of either OmpC or OmpF and their analogs was down in 92.1% of non-CP-CRE isolates. Although only 60.5% of non-CP-CRE encoded an ESBL or AmpC, respectively, the Check-Points assay does not detect chromosomally-encoded AmpCs present in the “SPACE” organisms (i.e., *Serratia*, *Citrobacter*, and *Enterobacter*), which accounted for 63.2% of non-CP-CRE isolates in this study. Overall, between carbapenemase and porin loss detection, we could account for mechanism of resistance in 95.2% (59/62) of CRE isolates in this study. The mechanism of resistance in unaccounted isolates may include efflux pump and target modification (36, 37).

The antibiotic susceptibility findings from this study are consistent with prior studies showing the underlying mechanism of carbapenem resistance predicts *in vitro* susceptibility to newly-developed β-lactam-β-lactamase inhibitor combinations, three of which (i.e., ceftazidime-avibactam, meropenem-vaborbactam, and ceftolozane-tazobactam) are FDA cleared and commercially available (12–14, 38). Susceptibility of CP-CRE isolates to ceftazidime-avibactam, meropenem-vaborbactam, and imipenem-relebactam was dependent on the molecular class of carbapenemase they encoded such that 100% of isolates encoding *bla*_KPC_ were susceptible to ceftazidime-avibactam, meropenem-vaborbactam and imipenem-relebactam; 100% of isolates encoding *bla*_OXA-48 like_ were susceptible to ceftazidime-avibactam but not to meropenem-vaborbactam and imipenem-relebactam; and 100% of isolates encoding metallo β-lactamases (i.e., *bla*_NDM_, *bla*_IMP_, and *bla*_VIM_) were non-susceptible to ceftazidime-avibactam, meropenem-vaborbactam, and imipenem-relebactam, excluding one *bla*_IMP_-positive isolate that was susceptible to imipenem alone. In non-CP-CRE isolates, 100%, 92.1%, and 89.5% were susceptible to ceftazidime-avibactam, meropenem-vaborbactam, and imipenem-relebactam, respectively. Susceptibility of non-CP-CRE isolates to imipenem-relebactam is consistent with findings by Livermore and colleagues but not by Lapuebla and colleagues (14, 39). This discrepancy could be due to extend of porin inactivation among non-CP-CRE isolates in different studies given that we showed isolates with resistance to imipenem-relebactam had nearly undetectable porins. Overall, our findings indicate nucleic acid testing for *bla*_KPC_, *bla*_OXA-48 like_, *bla*_NDM_, *bla*_IMP_, and *bla*_VIM_ is sufficient to distinguish between CP-CRE and non-CP-CRE and to accurately predict susceptibility to ceftazidime-avibactam, meropenem-vaborbactam, and imipenem-relebactam at our institution. Although not developed for treatment of CP-CRE, 100% of *S. marcescens* encoding *bla*_SME_ were susceptible to ceftolozane-tazobactam while the rest of CP-CRE were resistant or intermediate. Further, 31.6% of non-CP-CRE were susceptible to ceftolozane-tazobactam, however, genotypic cephalosporinase testing could not identify this group.

Although our findings are informative for management of patients with CRE infection, this study has several limitations. First, this was a single-center study. Given that patient population, medical management, and infection control practices vary between health systems, our findings might not be generalizable. Thus, a multicenter study in our geographic region is needed to confirm our findings. Second, despite including all CRE isolates at our institution over a 4-year period, the total number of CRE isolates was relatively small. However, this reflects natural epidemiology of CRE at our institution. Studies with large number of CRE isolates are needed to confirm our findings. Third, we compared the susceptibility of CRE isolates to four β-lactam combination drugs using three different susceptibility testing methods. Although our findings were consistent with prior reports, using a single susceptibility testing method for all four drugs might have allowed for more accurate comparison between drugs.

In conclusion, comprehensive phenotypic and genotypic characterization of CRE isolates longitudinally at a health system in Silicon Valley identified diverse resistance mechanisms including representation of all plasmid-encoded carbapenemases. On demand nucleic acid testing was able to accurately distinguish between CP-CRE and non-CP-CRE and predict *in vitro*susceptibility to ceftazidime-avibactam, meropenem-vaborbactam, and imipenem-relebactam.

## Acknowledgment

The study was supported by a research grant from Merck Inc to Niaz Banaei. We thank Robin Patel from the Mayo Clinic, Rochester, MN; Kyungwon Lee from Yonsei University College of Medicine, Seoul, South Korea; Andreas Wendel from Heinrich Heine University Düsseldorf, Germany and JMI laboratories, IA for providing bacterial isolates.

